## Supplementary Information for "Segment-Specific Optogenetic Stimulation in *Drosophila melanogaster* with Linear Arrays of Organic Light-Emitting Diodes"

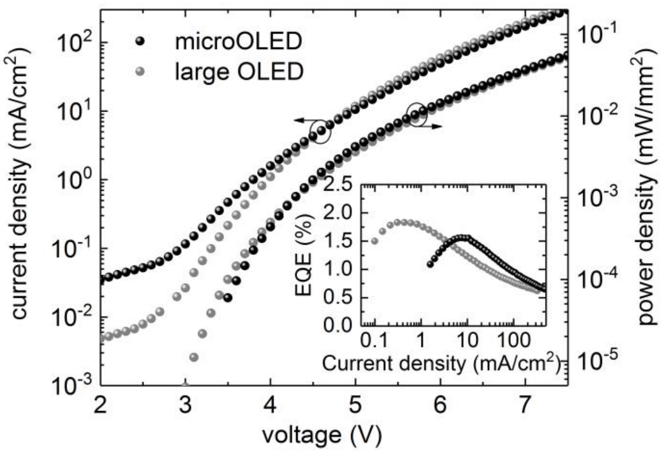


**Figure S1.** Current density and optical power density as a function of the voltage for large (4×4mm$^2$) and for microstructured OLEDs. Inset: External quantum efficiency as a function of the current density.

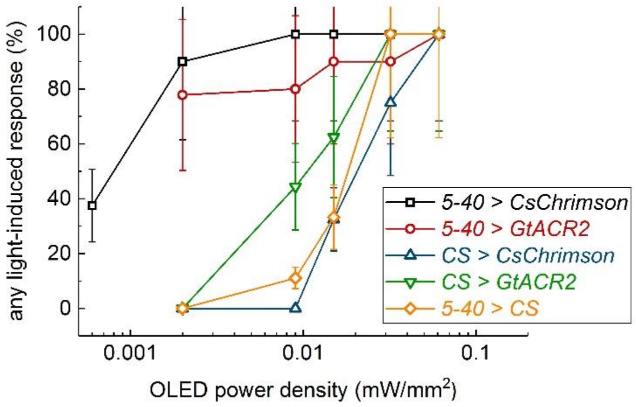


**Figure S2.** Dose-response curve for detecting any light-induced behaviour, e.g., twitching or stronger head movements, during exposure to blue light illumination from 4×4 mm$^2$ OLEDs. Control larvae showed a light-evoked behaviour for intensities of around 15 µW mm$^{-2}$ and higher. Data shows mean ± SEM; $n \geq 7$ larvae in each genotype.

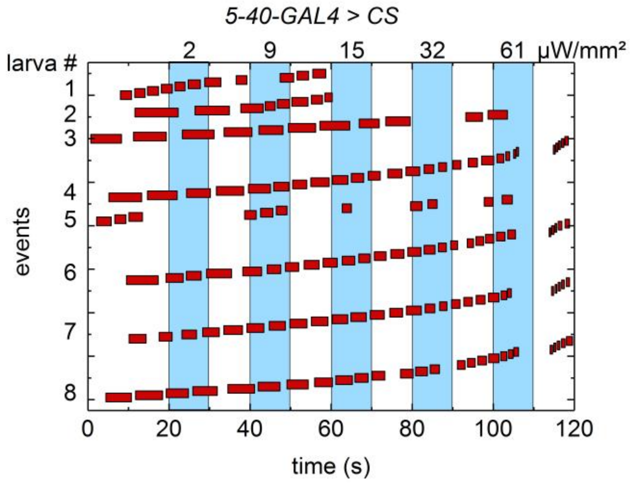


**Figure S3.** Wave duration of forward muscle contraction waves upon stimulation with a large OLED at different brightness levels (OLED power density as indicated in top axis) for 5-40-GAL4 > CS larvae. Different events are spread across the y-axis. Numbers label events from individual animals (*n* = 8 larvae). Muscle contraction waves significantly accelerated at power densities above 32 µW mm$^{-2}$ and were accompanied with strong head and tail movements (Supporting Video 3).

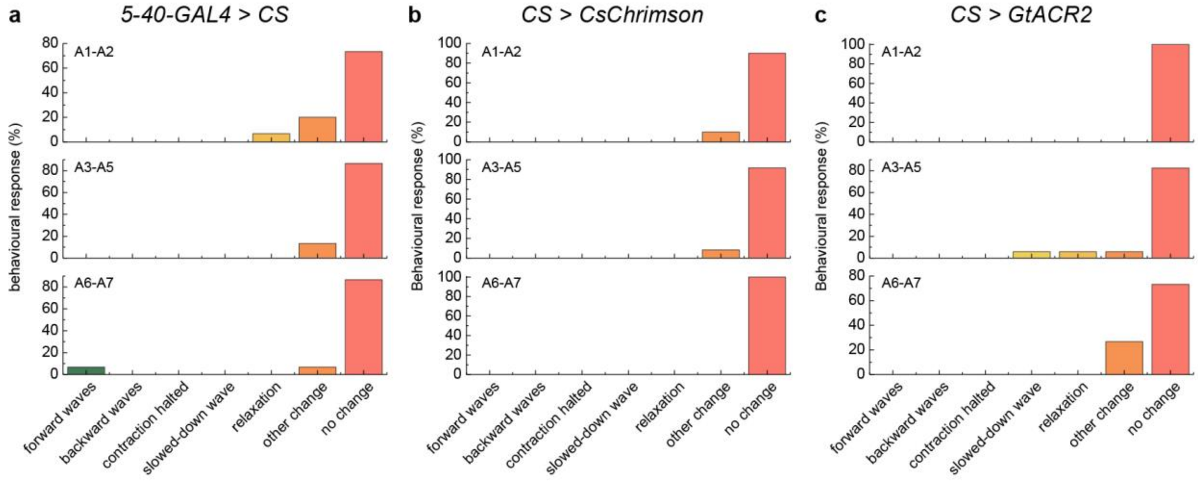


**Figure S4.** Behavioural response of controls upon local stimulation with three neighbouring OLED pixels delivering 15 µW mm$^{-2}$ for 10 s. Most larvae (> 70 % for each condition) showed no response. Some larvae showed a change in behaviour, however, this is attributed to random, non-light-induced changes. Importantly, no muscle contraction waves were evoked. *n* ≥ 4 larvae for each condition.

**Video 1.** Whole animal stimulation during forward crawling in a GtACR2-expressing larva using a large OLED pixel. Upon stimulation, relaxation of the larva or slow-down of muscle contraction waves were recorded. The 10 s-long periods during which the OLED is on are indicated in the video. OLED power density: 15 μW $mm^{-2}$.

**Video 2.** Dose-response upon whole animal stimulation of a CsChrimson-expressing larva using a large OLED pixel. The 3 s-long periods during which the OLED is on, the brightness, and the larval response are indicated in the video.

**Video 3.** Dose-response upon whole animal stimulation of a control larva using a large OLED pixel. The 10 s-long periods during which the OLED is on, the brightness, and the larval response are indicated in the video.

**Video 4.** Local stimulation of a CsChrimson-expressing larva in segment A5 using a microstructured OLED leads to a temporary stop of an incoming forward muscle contraction wave. Bottom: Location of the muscle contraction wave over time. The 3 s-long period during which the OLED is on is indicated in the video. OLED power density: 15 μW $mm^{-2}$.

**Video 5.** Switching the direction of locomotion of a CsChrimson-expressing larva by alternating stimulation of anterior and posterior segments with a microstructured OLED. Red lines indicate the current position of the muscle contraction wave. The 3 s-periods during which the OLED is on are indicated in the video. OLED power density: 15 μW $mm^{-2}$.
